# VARIOUS METHODS OF CHILLING AND HIGH-VOLTAGE ELECTRICAL STIMULATION APPLIED IN SUSTAINABLE BEEF PRODUCTION SYSTEMS

**DOI:** 10.1101/2020.10.01.321968

**Authors:** Joanna K. Banach, Ryszard Żywica, Paulius Matusevičius

## Abstract

Entrepreneurs implementing the concept of sustainable development of meat production are highly interested in combining various red meat chilling systems with quality-improving techniques. Therefore, we analyzed the impact of high-voltage electrical stimulation (HVES) and selected chilling methods on changes in quality characteristics and weight loss of beef. We also studied energy consumption based on the heat balance of chilling chambers during the fast chilling of varying amounts of raw material.

The HVES and the fast chilling method yielded positive economic (meat weight loss), technological (high quality, hot-boning), energetic (lower electricity consumption), and organizational effects (increased chilling speed, reducing of storage surfaces and expenditures for staff wages) compared to the slow and accelerated methods. Reaching the desired final temperature with an increased amount of chilled meat enabled a few-fold reduction in the total heat collected from the chambers and in meat weight loss. This allows recommending the above actions to be undertaken by entrepreneurs in the pursuit of sustainable meat production.

## INTRODUCTION

The meat industry has a high demand for electricity, with chilling systems (chilling compressors) consuming by far the most of this type of energy (Feliciano et al. 2014; Ferrarez et al. 2016). Given that the profitability of sales of Polish meat processing plants is at 1-2% and energy saving plays a vital role in the sustainable development of the meat industry, optimization of its consumption becomes a must (Pathare, Roskilly, & Jagtap 2019). Producers seek to reduce electricity costs and to increase energy efficiency in the meat industry’s innovative development, including the implementation of scientific research at the stage of cattle slaughter and the chilling process of beef carcasses/half-carcasses. Nevertheless, the investment and modernization measures can be undertaken after checking the extent of their impact on production economics and reducing their adverse impact on the environment in practice. Only this would allow the entrepreneurs to implement voluntary quality and environment management systems (Fritzson & Berntsson 2006a, b; Tanaka 2011; Alcázar-Ortega et al. 2012; Song, Wang & Ma 2019).

Today, the development of meat chilling techniques is heading towards systems that do not use extremely low temperatures, and yet allow for low meat weight losses. To improve the energy consumption and economy of beef production, slow methods are replaced by fast and shock methods. In practice, however, too slow or too fast chilling of carcasses may result in their deteriorated quality. Therefore, great caution should be exercised when choosing the right chilling method (Braden, 2013; Zhang et al. 2019). An adverse technological phenomenon that can occur during rapid meat chilling is the *cold shortening*. It happens when meat temperature drops below 10 °C at a pH value above 6.2, i.e. when the muscles are before or during the *rigor mortis* state. To prevent this phenomenon, meat producers use various physical methods to accelerate biochemical changes in meat in the slaughter / pre-chilling period. These include: tenderstretching (hanging carcasses by the Achilles tendon / pelvic bone), electrostimulation, or delayed chilling-conditioning (Banach & Żywica 2010; Kim et al., 2012; Beffin et al. 2018; Mikołajczak et al. 2019). Electrical stimulation (ES) is considered an ideal non-invasive solution in this case. However, its implementation in practice depends on the specifics and capabilities of a meat processing plant, including its production efficiency, mechanization of production processes as well as the use of processing power and surfaces (Swords et al. 2008; Savell 2012; Wojdalski, Dróżdż & Lipiński, 2010; Wojdalski & Niżnikowski, 2019).

Research trends in the field of combining various red meat cooling systems with available techniques to improve its quality are still investigated (Zhang et al. 2019). They are also in the focus of interest of entrepreneurs who are obliged to implement the concept of sustainable development of meat production (Pagan, Renouf, & Prasad 2002; Ramirez, Patel, & Blok, 2006; Pimentel et al. 2008; Song, Wang & Ma, 2019). Therefore, this study aimed to determine the impact of using an own-construction device for high-voltage electrical stimulation and selected chilling methods (slow, accelerated, fast) on changes in basic quality characteristics of beef, meat weight loss, and energy consumption of the chilling process based on the heat balance of chilling chambers in the process of fast cooling with varying amounts of raw material.

## MATERIAL AND METHODS

The research material consisted of carcasses / half-carcasses of meat cattle (heifers, bulls-age around 18 months, cows-age around 10 years) of the Polish Holstein-Friesian Black and White breed with an average carcass weight of approx. 200 kg. Stunning and post-stunning procedures were executed under industrial conditions following provisions of the Council Regulation (EC) No 1099/2009 of 24 September 2009 on the protection of animals at the time of killing. The study was carried out in a medium-sized meat processing plant with a capacity of the slaughter line reaching 100 carcasses per shift.

### High-voltage electrical stimulation

Approx. 40 min after stunning, carcasses was subjected to a high-voltage electrical stimulation (HVES) using a device designed by Żywica and Banach (2014), which won a bronze medal at the 45^th^ International Exhibition of Inventions and Innovations “Brussels Eureka 96” in Brussels. The HVES was performed with an alternate electrical current at U = 330V, f = 17Hz, rectangular shape of impulses, and pulse-duty factor 0.9 for 120 sec. Next, the carcasses were dissected into half-carcasses, flushed with water having a temp. of ca. 10°C, weighed, and transferred to chilling chambers.

### Chilling methods

The beef half-carcasses were chilled with the slow, accelerated, and fast two-phase methods, under the following conditions:

a. ***Slow method (delayed chilling):*** After leaving the slaughter line, the half-carcasses were transferred to a chilling chamber with air temp. of ca. +10°C, where they were kept for ca. 6h until the chamber had been filled up. Afterward, cooling aggregators were turned on and the half-carcasses were chilled with air having a temperature of ca. 2°C and humidity of ca. 90 % for 24h.
b. ***Accelerated method***: After transportation to the chilling chamber, the half-carcasses were chilled with air having a temperature of ca. 2°C and humidity of ca. 90 % for 24h.
c. ***Fast two-phase method:*** After leaving the slaughter line, the half-carcasses were transferred to a fast chilling chamber with air temp. of ca. −8°C and humidity of 95%, where they were kept for 2.5h. Afterward, they were transferred to an equalizing (transitory) chamber to equalize temperatures of their outer and inner layers (for 0.5-3h depending on the number of half-carcasses), with the fans switched off. Next, they were dissected into quarter-carcasses and transferred to a chilling tunnel with air temp. of 0-2°C and humidity of ca. 90 % for ca. 16-18h.

In all chilling chambers, the half-carcasses were moved in a continuous pendular motion with strong forced convection.

### Temperature measurements

Meat temperature measurements were performed on 180 half-carcasses (n=60 in each chilling method) over the chilling time using a TES 1310 TYPE-K spike thermometer (accuracy of ± 0.1°C), at three different localizations of each half-carcass: *semimembranosus* muscle, *longissimus dorsi* muscle, and around a shoulder blade, at a depth of ca. 7cm. The measurements were performed 1, 3.5, 8, and 24h after stunning.

### pH measurements

Measurements of pH of the stimulated (ES, n=15) and control (C, n=10) beef chilled with the slow, accelerated, and fast methods were performed 2/3, 2, 6, and 24h after stunning with a pH-meter (CP-411, electrode OSH 12-01, Elmetron, Poland) in three different localizations in *m. longissimus thoracis et lumborum* between 7 and 8^th^ rib. Before the measurements made for meat from a given experimental group (ES, C), the electrode was washed with distiller water and the pH-meter was calibrated.

### Energy balance

The heat balance of the chilling chambers used in the fast chilling method was based on the amount of heat necessary to be removed from the fast chilling chamber (area of 84 m^2^) and the chilling tunnel (area of 57 m^2^) during one chilling cycle. The measurements were carried out in the spring season for 8 working days (D1-D8), with varying amounts of carcasses (n=10 to n=144) in the chamber.

The total heat (Q) collected from the chilling chamber (Eq.1) and its components were calculated using the following formulas (Eq. 2-5):

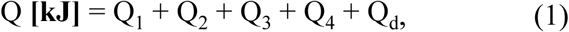

where; Q_1_ – heat removed form products (half-carcasses);

Q_2_ – heat transferring to the environment; the so-called transfer heat;

Q_3_ – ventilation heat;

Q_4_ – engine work heat;

Q_d_ – additional losses, in small chambers.

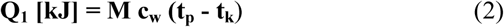

where: M – weight of chilled half-carcasses [kg],

c_w_ – specific heat capacity of half-carcasses [2.85 kJ/kg °C], [0.68 kcal/kg °C]

t_p_ – mean initial temperature of half-carcasses [°C]

t_k_ – mean final temperature of half-carcasses [°C]

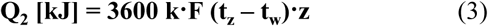

where: k – heat transfer coefficient of a building partition [0.44 W/m^2^ K]

F – size of a building partition [m^2^],

t_z_ – mean outer temperature of chamber environment [ºC],

t_w_ – mean inner temperature of the chamber [ºC],

z – heat transfer time [h].

The size of a building partition was assumed to be the size of the outer walls and ceiling of the chamber.

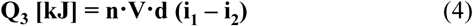

where: n – number of air exchanges in a chamber, for chilling chambers: n = 2 per day

V – cubic capacity of a chamber [m^3^],

d – mean air density in a chamber [kg / m^3^],

i_1_ – fresh air enthalpy [kJ / kg],

i_2_ – enthalpy of air in a chamber [kJ / kg].

Engine work heat (Q_4_) was assumed to be the electric energy consumed by engines installed in the chamber. It was calculated based on engine power and work time.

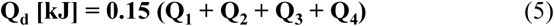

The heat of lighting and the heat of men’s work did not occur in the applied chilling technology. Data regarding: casing losses (Q_2_ – heat penetrating from the environment), total lighting, engine work, losses due to door opening, and fan work were derived from the ‘Operation and Maintenance Documentation’ available at the technical department of the audited plant.

### Weight losses of meat (WLM)

The effect of the fast chilling method on weight losses of stimulated meat (WLM) was determined based on the difference between the weight of half-carcasses before the chilling process (M_p_) and after the process (M_z_), and expressed in per cents (Eg. 6).

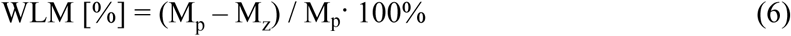

### Statistical analysis

The statistical analysis of the effects of chilling duration and method and of electrical stimulation on changes in meat temperature and pH was carried out using *Statistica* 13.1 software based on one-way analysis of variance (ANOVA, *p*<*0*.*01, p*<*0*.*05*).

## RESULTS AND DISCUSSION

### Effect of the chilling method on changes in meat temperature

Analyses carried out in the first part of the study were expected to allow understanding the mechanisms of the impact of various chilling methods on the basic meat quality attributes, expressed by changes in the temperature inside the meat and in meat pH, determining the rate of post-stunning changes, as well as quality and safety of beef during further storage or processing.

Results of meat temperature measurements demonstrated that the type of the chilling method (slow, accelerated, fast) had a significant effect on the rate of changes in muscle tissue temperature. Beef chilling with the slow method caused very slow but significant (*p*<*0*.*01*) changes in its temperature. The storage of beef half-carcasses in a chilling chamber for 2.5h at the elevated temperature (ca. +10°C/6h) contributed to meat temperature decrease by barely ca. 3°C, i.e. to ca. 36°C. In turn, after 8 and 24h of slow chilling, meat temperature dropped to ca. 25 and 5°C, respectively. Compared to the slow chilling method, slightly higher rate and significant (*p*<*0*.*01*) changes in meat temperature were demonstrated during accelerated chilling. As soon as after 2.5h of chilling, meat temperature decreased by ca. 13°C and reached ca. 26 °C. After 8h and 25h after stunning, it reached ca. 12°C and 4°C, respectively (Table 1). The above results corroborate literature data indicating that achieving this rate of changes in meat temperature in the first hours of post-stunning slow and accelerated chilling can pose the risk of microbiological contamination caused by the rapid proliferation of microorganisms on the surface of half-carcasses (Żywica, 2010). In contrast, the rate of changes in meat temperature was consistent with the ‘10/10 rule of thumb’, indicating that meat chilling to a temperature not lower than 10 °C within 10 h post-slaughter allows avoiding the ‘cold shortening’ phenomenon. The effect of counteracting this adverse phenomenon can be intensified by delayed/slow chilling, which involves keeping the half-carcasses outside the chilling room for some time (Smulders, Toldra’, Flores, & Prieto, 1992) or under either elevated temperatures, i.e., 10-16 °C / 3-12h (White, O’Sullivan, Troy & O’Neill 2006; Luo et al., 2008; Prado & de Felicio, 2010; Pflanzer, Gomes & De Felicio, 2019) or lower temperatures, i.e., 5-6 °C/24h (Janz et al., 2004).

**Table 1.** Temperature changes (mean values ± standard deviation) during beef chilling with the slow (SM), accelerated (AM), and fast (FM) methods

| Experimental group |  | Chilling methods |  |  | Significance |  |  |
| --- | --- | --- | --- | --- | --- | --- | --- |
|  |  | SM (n=60) | AM (n=60) | FM (n=60) |  |  |  |
| Time after stunning [h] | 1 | 38.8 ± 0.69<sup>a</sup> | 38.6 ± 0.57<sup>a</sup> | 38.2 ± 1.04<sup>a</sup> | NS | NS | NS |
| | 3.5 | 35.7 ± 1.15<sup>b</sup> | 25.7 ± 3.18<sup>b</sup> | 18.1 ± 1.57<sup>b</sup> | ** | ** | ** |
| | 8 | 25.3 ± 2.22<sup>c</sup> | 11.9 ± 2.58<sup>c</sup> | 10.3 ± 1.68<sup>c</sup> | ** | NS | ** |
| | 24 | 4.8 ± 0.45<sup>d</sup> | 3.8 ± 0.79<sup>d</sup> | 1.4 ± 0.95<sup>d</sup> | ** | ** | ** |

a, b, c, d – mean in columns with different letters differ significantly at $p < 0.01$ ;
\*\* - significance level $p < 0.01$ ; NS - non significant

The chilling of beef half-carcasses with the fast two-phase method had a strong effect on meat temperature decrease compared to the slow and accelerated methods. Keeping the half-carcasses in the chilling chamber for 2.5h caused its temperature to drop from ca. 38°C to ca. 18 °C. During further meat keeping in a chilling tunnel, its temperature dropped to ca. 10°C after 8h and to 1.4°C after 24h. The final temperature of meat chilled with the fast two-phase method was lower than of the meat chilled with the slow (ca. 5°C) and accelerated (ca. 4°C) methods. The statistical analysis of the effects of chilling methods on changes in meat temperature also demonstrated significant (*p*<*0*.*01*) differences between mean meat temperatures recorded after 3.5 and 24h of chilling with SM, AM, and FM as well as between mean meat temperatures measured after 8h of chilling with SM/AM and FM/SM (Table 1).

### Effect of chilling method and electrical stimulation on changes in meat pH

In addition to using a rapid cooling system, companies striving for sustainable beef production can employ various methods to model meat quality characteristics. Methods used in practice to naturally accelerate post-slaughter transformations in beef include electrical stimulation, both the low-voltage (ESNN, up to 100V) and the high-voltage one (HVES, from 100 to 3000V). The magnitude of their effects on biochemical processes in meat is determined by: animal type, process onset time, and choice of electrical current parameters (frequency, impulse shape, voltage type, etc.). Due to the possibility of exciting muscles through both the nervous system and muscle fiber conductivity, the HVES is considered the most effective method for meat quality improvement (Żywica, 2010). For this reason, the second part of this study aimed to determine the effect of high-voltage electrical stimulation of beef half-carcasses with an own-construction device (Żywica & Banach 2014) and their chilling with three methods: slow, accelerated, and fast two-phase, on the rate of changes in meat pH, being its main quality attribute (Contreras-Castillo et al. 2016; Banach et al. 2018). This complex approach was expected to allow identifying the optimal variant of meat production, ensure its high quality and reduce its harmful effect on environment.

Results of pH measurements of the *longissimus dorsi* muscle of ES and C heifers, cows, and bulls chilled with the slow method showed that HVES significantly (*p*<*0*.*001*) accelerated the post-stunning transformations during 24-h chilling. The highest and significant differences in pH values were obtained after 2 and 6 h of chilling. The pH_2h_ value of ES heifer, cow, and bull meat ranged from 5.98 to 6.10 and was lower by ca. 0.9 units compared to the pH_2h_ value of control samples. After 6h post-stunning, the pH value of ES meat was still significantly lower (by 0.7-0.8 units, *p*< *0*.*001*) compared to the pH values of meat of non-stimulated heifers, bulls, and cows that reached 6.42, 6.54, and 6.63, respectively. The final pH values (pH_24h_) of both ES and C meat were similar and reached before 5.7; however, the pH_24h_ values of ES and C meat of heifers and cows differed significantly at *p*<*0*.*05* (Table 2).

**Table 2.** Changes in the pH value 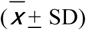 of the *longissimus dorsi* muscle of control (C) and electrically-stimulated (ES) heifers, cows, and bulls chilled with the slow method

| Experimental group |  | Heifers |  |  | Cows |  |  | Bulls |  |  |
| --- | --- | --- | --- | --- | --- | --- | --- | --- | --- | --- |
|  |  | ES (n=15) | C (n=10) | Significance | ES (n=15) | C (n=10) | Significance | ES (n=15) | C (n=10) | Significance |
| Time after stunning [h] | 2/3 | 6.95 ± 0.10<sup>a</sup> | 7.02 ± 0.09<sup>a</sup> | NS | 6.98 ± 0.11<sup>a</sup> | 7.00 ± 0.06<sup>a</sup> | NS | 6.86 ± 0.11<sup>a</sup> | 6.91 ± 0.08<sup>a</sup> | NS |
| | 2 | 5.98 ± 0.05<sup>b</sup> | 6.61 ± 0.05<sup>b</sup> | *** | 6.10 ± 0.08<sup>b</sup> | 6.87 ± 0.04<sup>b</sup> | *** | 6.03 ± 0.09<sup>b</sup> | 6.66 ± 0.02<sup>b</sup> | *** |
| | 6 | 5.75 ± 0.04<sup>c</sup> | 6.42 ± 0.03<sup>c</sup> | *** | 5.79 ± 0.07<sup>c</sup> | 6.63 ± 0.02<sup>c</sup> | *** | 5.82 ± 0.10<sup>cC</sup> | 6.54 ± 0.05<sup>c</sup> | *** |
| | 24 | 5.64 ± 0.05<sup>d</sup> | 5.73 ± 0.06<sup>d</sup> | * | 5.69 ± 0.03<sup>d</sup> | 5.75 ± 0.05<sup>d</sup> | * | 5.70 ± 0.12<sup>Dd</sup> | 5.78 ± 0.03<sup>d</sup> | NS |

<sup>a-d</sup> – means in the columns with different subscripts are significantly different, $P < 0.01$
\*\*\* - significance level $P < 0.001$ ; \*\* - significance level $P < 0.01$ ; \* - significance level $P < 0.05$ ; NS - non significant

The analysis of meat temperature recorded after ca. 3.5h of chilling (Table 1) and pH value of ES meat measured 2h post-stunning allows concluding that the use of HVES enables the hot-boning of meat as soon as 2h after stunning without the fear of its cold shortening during consecutive ripening. In also allows for eliminating the phase of meat retention in the chilling chamber (2.5h/ ca.10°C) and increasing production surface (Pisula & Tyburcy 1996, Simmons et al. 2006).

The analysis of pH values of meat chilled with the accelerated method (AM) demonstrated the rate of changes in pH values of C meat over the chilling period was the same (did not differ significantly) as in the slow method. Therefore, these changes were not presented in Table 2. In turn, the pH values of ES meat of heifers, cows, and bulls were insignificantly higher than in the slow chilling method (SM). The pH values measured in ES meat 2, 6, and 24h post-stunning fitted within the following ranges: pH_2h_ = 6.01-6.06, pH_6h_ = 5.73-5.87, and pH_24h_ = 5.6-5.7. The gender of animals had no significant effect on the final pH values (pH_24h_) of ES meat. However, the pH_24h_ value of bulls was higher and more differentiated (5.73±0.17) compared to the pH_24h_ value of heifers (5.60±0.05) and cows (5.65±0.04) -Table 3. Higher and more differentiated pH values of meat of the stimulated bulls proved that bulls are more susceptible to the pre-slaughter stress than heifers and cows (Żywica 2010). Negligible differences in the rate of pH changes between ES and control meat indicate that the AM chilling method is more rational and efficient than the SM, from the practical point of view.

**Table 3.**
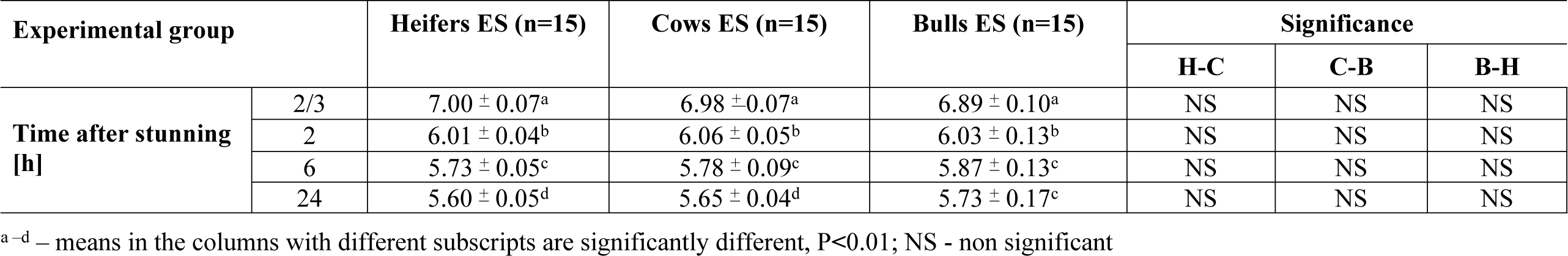
Changes in the pH value 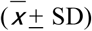 of the *longissimus dorsi* muscle of control (C) and electrically-stimulated (ES) heifers (H), cows (C), and bulls (B) chilled with the accelerated methods

| Experimental group |  | Heifers ES (n=15) | Cows ES (n=15) | Bulls ES (n=15) | Significance |  |  |
| --- | --- | --- | --- | --- | --- | --- | --- |
|  |  |  |  |  | H-C | C-B | B-H |
| Time after stunning [h] | 2/3 | 7.00 ± 0.07<sup>a</sup> | 6.98 ± 0.07<sup>a</sup> | 6.89 ± 0.10<sup>a</sup> | NS | NS | NS |
| | 2 | 6.01 ± 0.04<sup>b</sup> | 6.06 ± 0.05<sup>b</sup> | 6.03 ± 0.13<sup>b</sup> | NS | NS | NS |
| | 6 | 5.73 ± 0.05<sup>c</sup> | 5.78 ± 0.09<sup>c</sup> | 5.87 ± 0.13<sup>c</sup> | NS | NS | NS |
| | 24 | 5.60 ± 0.05<sup>d</sup> | 5.65 ± 0.04<sup>d</sup> | 5.73 ± 0.17<sup>c</sup> | NS | NS | NS |

<sup>a-d</sup> – means in the columns with different subscripts are significantly different, $P < 0.01$ ; NS - non significant

The use of lower temperatures during the fast chilling of ES meat (−8°C/2.5h, 1-2°C/ca. 16-18h) than in the accelerated and slow methods contributed to the slower post-stunning biochemical changes of meat and, consequently, to pH values higher by ca. 0.3 units at particular measuring points. While the temperature of stimulated meat chilled with the fast method reached 18 and 10°C after 2 and 8h since stunning, its pH_2h_ was at ca. 6.2 and 6.3, and its pH_6h_ was at 5.95 (Table 4). This means that the coupling of HVES treatment performed with the own-construction device and the fast chilling method allows achieving a very rapid temperature drop before the onset of *rigor mortis* and producing high-quality meat without the fear of its cold shortening (Li et al., 2012; Pinto Neto et al., 2013; Devine et al., 2014; Sikes et al. 2017). In the case of control meat, at pH_2h_ = 6.7-7.8 and pH_6h_ = 6.5-6.6, the biochemical processes are still in progress, and meat is still before the *rigor mortis* state; therefore, its rapid chilling without ES is not recommendable (Strydom, Frylinck & Smith 2005).

**Table 4.** Changes in the pH value 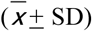 of the *longissimus dorsi* muscle of control (C) and electrically-stimulated (ES) heifers (H), cows (C), and bulls (B) chilled with the fast method

| Experimental group |  | Heifers (n=15) |  |  | Cows (n=15) |  |  | Bulls (n=15) |  |  |
| --- | --- | --- | --- | --- | --- | --- | --- | --- | --- | --- |
|  |  | ES | C | Significance | ES | C | Significance | ES | C | Significance |
| Time after stunning [h] | 2/3 | 6.87 ± 0.11<sup>a</sup> | 6.91 ± 0.05<sup>a</sup> | NS | 7.02 ± 0.06<sup>a</sup> | 7.00 ± 0.03<sup>a</sup> | NS | 6.92 ± 0.10<sup>a</sup> | 6.95 ± 0.09<sup>a</sup> | NS |
| | 2 | 6.24 ± 0.08<sup>b</sup> | 6.67 ± 0.02<sup>b</sup> | *** | 6.29 ± 0.11<sup>b</sup> | 6.80 ± 0.04<sup>b</sup> | *** | 6.35 ± 0.07<sup>b</sup> | 6.71 ± 0.06<sup>b</sup> | ** |
| | 6 | 5.95 ± 0.12<sup>c</sup> | 6.50 ± 0.07<sup>c</sup> | *** | 5.92 ± 0.09<sup>c</sup> | 6.65 ± 0.04<sup>c</sup> | *** | 5.95 ± 0.11<sup>c</sup> | 6.53 ± 0.07<sup>c</sup> | *** |
| | 24 | 5.69 ± 0.06<sup>d</sup> | 5.78 ± 0.06<sup>d</sup> | * | 5.74 ± 0.10<sup>d</sup> | 5.72 ± 0.08<sup>d</sup> | NS | 5.70 ± 0.05<sup>d</sup> | 5.75 ± 0.04<sup>d</sup> | NS |

<sup>a-d</sup> – means in the columns with different subscripts are significantly different, $p < 0.01$
\*\*\* - significance level $p < 0.001$ ; \*\* - significance level $p < 0.01$ ; \* - significance level $p < 0.05$ ; NS - non significant

Given the results presented above, it has been concluded that the coupled use of HEVS performed using an own-construction device and a fast chilling method is justified considering both the economic and quality concerns.

### Energy consumption of the meat chilling with the fast method

The rational management of electrical energy in the production process entails not only accomplishing economic effectiveness (increasing savings) but also reducing the energy consumption of technological processes and facilities, which determine the magnitude of effects of technical and man activities on the natural environment (Pagan, Renouf, & Prasad 2002; Pelletier 2008). The chilling systems are the largest electricity consumers in the meat industry – they account for 50-93% of the total electric energy consumed in a typical slaughterhouse (Nunes, Silva, Andrade 2011; Feliciano et al. 2014). Therefore, as part of internal cost control / production savings and preparation for the energy audit (Munguia et al. 2016), entrepreneurs (with individual production specifics) should keep records of the total consumption of energy carriers. Due to the above facts, the indicator of the degree of implementation of the sustainable energy policy by the meat industry in practice is most often expressed in unitary consumption of electric energy, determined by the heat balance of chilling chambers (Ramírez, Patel, & Blok, 2006; Tasmania, 2009; Wojdalski, Dróżdż & Powęzka, 2009; Wojdalski et al. 2013).

Results of measurements and calculations of the heat balance of the fast chilling chamber (FChCh) and the chilling tunnel (ChT) demonstrated that the heat of penetration, lighting heat, and engine work heat in these rooms had constant values (FChCh=33.94, ChT=23.26). The total amount of heat necessary to be removed from the chambers depended on the duration of their work and the amount of raw material being chilled (half-carcasses, quarter-carcasses). In turn, the weight and type of chilled raw material, as well as the final temperature of chilled meat (0-2°C) determined the amount of heat that had to be removed from it (Fontana et al. 1999; Kaliszan et al. 2005). Apart from the mentioned factors, the total amount of heat depended on additional heat losses associated with door opening (10-15%) and heat emitted by ventilation fans (20%) – Table 5.

**Table 5.** Heat balance of the fast chilling chamber (FChCh) and chilling tunnel (ChT), and changes in weight losses of meat during fast chilling for 8 days (D1-D8)

| No | Specification |  | Unit | D1<br>(n=10) | D2<br>(n=21) | D3<br>(n=26) | D4<br>(n=42) | D5<br>(n=50) | D6<br>(n=59) | D7<br>(n=68) | D8<br>(n=114) |
| --- | --- | --- | --- | --- | --- | --- | --- | --- | --- | --- | --- |
| 1 | Casing losses ($Q_2+Q_4$) | FChCh | kJ/h | 33.94 | | | | | | | |
|  |  | ChT |  | 23.26 |  |  |  |  |  |  |  |
| 2 | Work time | FChCh | h | 3.5 | 4.5 | 5.0 | 6.5 | 8.0 | 7.5 | 7.5 | 8.5 |
|  |  | ChT |  | 18.0 | 18.0 | 17.5 | 16.0 | 14.5 | 15.0 | 15.0 | 14.0 |
| 3 | 1×2 | FChCh | kJ | 118.79 | 152.73 | 169.70 | 220.61 | 271.52 | 254.55 | 254.55 | 288,49 |
|  |  | TCh |  | 418.68 | 418.68 | 407.05 | 372.16 | 337.27 | 348.90 | 348,9 | 325.64 |
| 4 | Heat removed from half-carcass ($Q_1$) | FChCh | kJ | 178.90 | 325.72 | 347.66 | 600.64 | 609.11 | 616.41 | 622,73 | 1009.70 |
|  |  | ChT |  | 61.90 | 147.56 | 158.27 | 264.99 | 291.23 | 322.63 | 386.56 | 631.84 |
| 5 | 3+4 | FChCh | kJ | 297.69 | 478.45 | 517.36 | 821.25 | 880.63 | 870.96 | 877.28 | 1298.19 |
|  |  | ChT |  | 480.58 | 566.24 | 565.32 | 637.09 | 628.50 | 671.53 | 735.46 | 957.48 |
| 6 | Losses due to door opening ($Q_d$) | FChCh | % | 10 | | | | | | | |
|  |  | ChT |  | 15 |  |  |  |  |  |  |  |
| 7 | Heat emitted by ventilation fans ($Q_3$) | | % | 20 | | | | | | | |
| 8 | Total heat removed from: | FChCh | kJ | 391.54 | 632.99 | 684.57 | 1084.05 | 1174.82 | 1145.25 | 1158.00 | 1717.23 |
|  |  | ChT |  | 660.52 | 775.42 | 797.18 | 879.14 | 898.48 | 966.57 | 1014.83 | 1331.37 |
| 9 | Heat removed per carcass<br>Total heat removed from<br>(5×6×7) | FChCh | kJ/carcass | 39.15 | 30.14 | 26.33 | 25.81 | 23.50 | 19.41 | 17.03 | 15.06 |
|  |  | ChT |  | 60.05 | 36.40 | 30.67 | 20.93 | 17.97 | 16.38 | 14.92 | 11.68 |
| 10 | Total heat removed from FChCh and ChT per carcass |  | kJ/carcass | 99.20 | 66.54 | 57.00 | 46.74 | 41.47 | 35.79 | 31.95 | 26.74 |
| 11 | Initial weight |  | kg | 2985 | 5569 | 6616 | 10331 | 9866 | 10553 | 12072 | 31437 |
| 12 | Final weight |  | kg | 2842 | 5369 | 6411 | 9969 | 9639 | 10300 | 11746 | 30620 |
| 13 | Weight losses of meat |  | % | 4.8 | 3.1 | 3.1 | 3.5 | 2.3 | 2.4 | 2.7 | 2.6 |
| 14 | Average weight loss of meat (n=390) |  | % | 2.8 |  |  |  |  |  |  |  |

$Q_1$ – heat removed form products (half-carcasses); $Q_2$ – heat transferring to the environment; the so-called transfer heat; $Q_3$ – ventilation heat; $Q_4$ – engine work heat; $Q_d$ – additional losses, in small chambers; FChCh - fast chilling chamber; ChT - chilling tunnel

Results of measurements and calculations demonstrated that the total heat needed to be removed from chilling chambers increased along with the increasing number of carcasses intended for chilling (10-114 animals). However, ca. 11-fold increase in the number of chilled carcasses caused over 4-fold increase (391.54 – 1717.23 kJ) in the fast chilling chamber (FChCh), and a 2-fold increase (660.52 – 1331.37 J) in the chilling tunnel (ChT) in the amount of heat that had to be removed from these rooms (Table 5). This indicates that the heat load of the chilling chamber (production of the amount of chill and electrical energy consumption) increases along with the increasing load of products placed in it within a day, with the increasing specific heat of the chilled product, and with the increasing difference of temperatures (Nunes, Silva & Andrade, 2011). Keeping the temperature lower than the ambient temperature in cold rooms requires also discharging heat energy from them, which entails higher losses (longer work of compressors) that will generate greater costs of energy consumption. The amount of heat that needs to be removed from chilling chambers is also strongly affected by the ambient temperature related to the season of the year. According to Nerynga et al. (1990), greater by ca. 32% electric energy consumption per chilled product unit is recorded in the spring than in the winter. The higher temperature around the chilling chamber contributes to, e.g., slower cooling of half-carcasses after slaughter, the higher temperature in the chamber before the chilling process, and a higher value of the transfer heat.

Compared to the calculations of the total heat capacity, opposite results were derived from calculations per carcass (unitary total heat capacity). With an over 9-fold increase in the number of carcasses, it was necessary to remove ca. 3-fold lower heat capacity unit from the fast chilling chamber (39.15-15.06 kJ/animal) and over 5-fold lower heat capacity unit from the chilling tunnel (60.05-11.68 kJ/animal). This was due to the high contribution of heat losses from casings (heat transfer, lighting, engine work, door opening) in the total amount of heat needed to be removed from the chambers. This, in turn, is associated with chamber work time. Żywica, Banach and Gornowicz (2008), obtained the unitary electric energy consumption by compressors at 19 MJ/t by chilling 10,320 kg of meat with the fast two-phase method. In contrast, the energy consumption decreased almost twice (10.27 MJ/t) when they used the shock chilling method. By decreasing the weight of the chilled meat (6182 kg), they reduced energy consumption to 15 and 4.03 MJ/t, respectively. In turn, Nunes, Silva and Andrade (2011) demonstrated that the meat industry is characterized by higher unitary consumption of electric energy 1208 kWh/t raw material than the other industries in Portugal. To achieve energy savings of around 17%, the Authors recommend it would be required, among other things, to improve the power management procedures and also of the activities, so that the chambers wouldn’t stay running at partial load.

Considering that one chilling cycle in a fast chilling chamber lasts 2.5h, its prolongation by 1h with carcass number of 10, by 2-3.5h with carcass numbers of 22 and 26, and by 6.5h with carcass number of 114 was due to the stable time of cooling the chambers before they were filled with half-carcasses (regardless of the amount of chilled material) and the stable chilling time of one half-carcass in the chamber. Besides, the temperature decrease by ca. 20°C in the fast chilling chamber and by ca. 18°C in the tunnel (14 to 18h, Table 1) influenced the total unitary amount of heat that had to be removed from both types of chambers (FChCh and ChT). During chilling, the total heat capacity per carcass was 99.20 kJ with 10 carcasses, and ca. 4-fold lower (26.74 kJ) with 114 carcasses (Table. 5).

An indispensable side effect of the meat chilling process are its weight losses. They generate economic losses, the magnitude of which depends on the chilling method type. According to Brown et al. (2009), the cost incurred by small UK plants due to meat weight losses was higher than the cost of electricity. Therefore, quick chilling methods are recommended (Zhang eta al. 2019) to reduce meat weight loss and get the best efficiency of the chilling process (shortening the time, ensuring meat safety and quality).

Results of determinations of weight losses of meat (WLM) caused by water evaporation from meat surface during its chilling with the fast two-phase method demonstrate large differences between particular days of measurements assuming various numbers of animal carcasses to be chilled (Table 1). The highest WLM (4.8%) was obtained during chilling 10 beef carcasses, presumably due to the highest volume of water evaporated from half-carcasses in the fast chilling chamber. On days 2, 3, and 4 of measurements, when the number of chilled carcasses was 21-42, the WLM ranged from 3.1 to 3.5%. The results of measurements allow concluding that smaller WLM were recorded (2.3-2.6%) with carcasses having a higher meat content, more densely packed, and touching one another in the chamber than with the lower number of chilled carcasses (Table 5). According to Klettner (1996), the weight loss of meat caused by its chilling with the fast one-phase method was at 1.6% and was higher by ca. 0.3% from the WLM recorded during the shock chilling process (from –3 to –5^°^C/2h, 0^°^C/16–22h). However, the carcasses intended for chilling should not be moist because of the possibility of meat surface freezing during chilling with this method. As reported by Żywica, Banach and Gornowicz (2008), WLM obtained using the shock and fast chilling methods reached 2.2 and 3.6%, respectively.

## CONCLUSIONS

1. The use of HVES and the fast chilling method at the slaughter line enables producing high-quality meat, reducing expenditures related to electric energy consumption, and increasing work effectiveness of cooling appliances compared to the slow and accelerated chilling methods. The coupling of these techniques is recommended for both investment and modernization undertakings in pursuit of sustainable meat production.
2. The intensive chilling of beef to ca. 20°C within 2.5h and the break in the chilling process until temperature equalization between its outer and inner layers allow the producers for getting the optimal rate of pH changes, rational quality management and changes in production organization, mainly through increasing chilling speed and reducing storage surface and expenditures for staff wages.
3. The heat balance of fast chilling chambers (FChCh+ChT) demonstrated several times higher unit amount of total heat needed to be removed and also greater meat weight losses when the final desired meat temperature was achieved with the lowest than with highest number of chilled carcasses.
4. The assessment of quality and energetic attributes of beef production will allow determining practical and available energy efficiency measures for chilling equipment in the meat plant under examination that will improve its image and reduce its harmful effect on environment.

